# Elephant seal cells export iron and form lipid droplets to evade lipid peroxidation

**DOI:** 10.64898/2026.08.10.744012

**Authors:** Kaitlin N Allen, Elizabeth R Piotrowski, Diana D Moreno-Santillán, Alexander L Li, Diamond Luong, Violet E Foley, Caroline del Real, José Pablo Vázquez-Medina

**Affiliations:** Department of Integrative Biology, University of California, Berkeley, Berkeley, CA; Department of Biology, Woods Hole Oceanographic Institution, Woods Hole, MA; University of Maryland Center for Environmental Science, Baltimore, MD; Department of Biology, University of San Francisco, San Francisco, CA; Department of Molecular and Cell Biology, University of California, Berkeley, Berkeley, CA; Department of Biochemistry and Molecular Biology, Brown University, Providence, RI

**Keywords:** Endothelial cells, ferroptosis, ACSL3, FPN1, ischemia/reperfusion

## Abstract

Elephant seals are remarkable breath-hold divers, capable of remaining submerged for up to two hours during diving bouts. These dives entail repeated, extreme hypoxia/reoxygenation events that would induce severe lipid peroxidation and tissue dysfunction in most species. Here, we show that primary vascular endothelial cells derived from elephant seals resist oxidant-induced lipid peroxidation and cell death without relying on canonical lipid hydroperoxide scavenging. Comparative transcriptomic profiling across seal, human, and sheep cells identified transcriptional signatures of lipid metabolism and iron handling as uniquely regulated in seal cells following hydroperoxide exposure. Mechanistically, seal cells exhibit high constitutive levels of acyl-CoA synthetase long-chain family member 3 (ACSL3) alongside robust, seal-specific induction of the iron exporter ferroportin-1 (FPN1). Comparative lipidomic profiling revealed that seal cells are enriched in triglycerides and monounsaturated fatty acids. ACSL3 silencing and FPN1 inhibition each partially sensitized seal cells to lipid peroxidation, identifying lipid droplet biogenesis and active iron export as complementary but non-redundant axes of protection. Together, these data reveal a protective strategy based on lipid sequestration and iron export rather than hydroperoxide reduction, with implications for ischemia-reperfusion injury in non-diving mammals.

**New and Noteworthy:** Deep-diving elephant seals repeatedly experience extreme hypoxia-reoxygenation events without incurring vascular injury. Here we show that vascular endothelial cells derived from elephant seals evade lipid peroxidation using a dual-defense axis that combines lipid droplet biogenesis to sequester vulnerable lipids with rapid iron export. This study reveals naturally evolved cellular mechanisms that confer protection against vascular ischemia/reperfusion.

## Introduction

Elephant seals are premier divers that spend up to 90% of their at-sea time submerged, performing prolonged, repeated dives (1). Diving seals utilize a suite of cardiovascular adjustments, including selective vasoconstriction, to modulate tissue oxygen consumption (2–5). Additionally, seals rely heavily on the heme-based respiratory pigments hemoglobin and myoglobin to bolster body oxygen stores (6). Despite these adaptations, elephant seals regularly experience arterial PO_2_ values as low as 12 mm Hg, corresponding to an arterial sO_2_ of ∼8% while diving. These extreme hypoxemic events are followed by rapid reoxygenation during brief eupneic surface intervals (7, 8). In humans, arterial sO_2_ values below 70% are a life-threatening emergency, and localized fluctuations in oxygen availability on par with seal values occur only during adverse cardiovascular events such as myocardial infarction and ischemic stroke (9, 10).

Hypoxia-reoxygenation events trigger oxidant generation, disrupting redox homeostasis and causing oxidative damage (10). Concurrently, iron accumulates during hypoxia. Upon reoxygenation, ferrous iron reacts with molecular oxygen, driving the peroxidation of membrane phospholipids (11). Excessive lipid peroxidation – a hallmark of ischemic disease – ultimately triggers ferroptosis, a form of regulated, iron-dependent cell death (12, 13). To counteract lipid peroxidation, glutathione (GSH)-dependent peroxidases, notably GPX4, reduce toxic phospholipid and short-chain hydroperoxides to their corresponding alcohols (13, 14). While abundant evidence links GSH metabolism to oxidative stress resistance in marine mammals, the precise mechanisms underlying how seal cells evade lipid peroxidation remain uninvestigated (15–19).

Vascular oxidative damage disrupts endothelial barrier integrity and impairs vasoregulation (20–23). Therefore, investigating how elephant seals naturally withstand vascular oxidative stress offers an opportunity to identify novel mechanisms of vascular protection. Here, we established primary arterial endothelial cell cultures from elephant seal arteries to study how these extreme diving mammals naturally cope with vascular oxidative stress. We found that, in contrast to vascular endothelial cells from humans or sheep, seal cells can evade lipid peroxidation. Using comparative transcriptomics, we identified gene expression signatures associated with iron handling and lipid remodeling as potential cytoprotective mechanisms in seal cells. Further experiments using comparative lipidomics and pharmacological and genetic tools revealed that seal cells evade lipid peroxidation by forming lipid droplets and actively exporting iron. In sum, our results show that extreme-diving mammals use a multifaceted, coordinated strategy to avoid vascular oxidative stress.

## Methods

### Tissue acquisition, primary cell isolation, and culture

Northern elephant seal (*Mirounga angustirostris*) and human placental arterial endothelial cells were derived as described in (24). Seal tissue was obtained under NMFS permit # 19108 (PI: Daniel Costa, UC Santa Cruz). Seal cells were isolated and cultured under NMFS permit # 22479 (PI: José Pablo Vázquez-Medina, UC Berkeley). Sheep *(Ovis aries)* placentae were obtained from the UC Davis School of Veterinary Medicine and cultured in commercial endothelial cell medium (Sciencell Research Laboratories). Confluent cultures were cryopreserved at passage 1. Seal and human endothelial cell cultures were previously characterized (24). The endothelial phenotype of sheep cell cultures was confirmed by immunostaining for VE-cadherin (CD144; ThermoFisher cat. # PA5-17401; RRID:AB_10987564; 1:100) **(Fig S1)**. All experiments were conducted in cells at passage 4-7 cultured in commercial endothelial cell medium, and species were passage-matched within each experiment.

### Hydroperoxide treatment and hypoxia-reoxygenation exposure

Confluent cells were exposed to increasing concentrations of *tert*-Butyl hydroperoxide (*t-*BOOH; 0, 100, 200, 300 µM) for up to 2 h. Live cells were used for cell viability/cytotoxicity and imaging experiments or harvested for RNA-seq, lipidomics, RT-qPCR, western blot, or activity assays. For hypoxia-reoxygenation experiments, cells were incubated in an InVivO_2_ physiological cell culture workstation (Baker Ruskinn) for 1 h at 0.5% O_2_, 5% CO_2_, and 37°C. The cell culture medium was deoxygenated in the workstation overnight to ensure the rapid onset of hypoxia (24, 25). The medium in each dish was replaced with deoxygenated medium when cells were introduced to the station. Reoxygenation was achieved by removing cells from the chamber and briefly opening dish lids in a biosafety cabinet in room air (21% O_2_), followed by 30 min in a 5% CO_2_ incubator at 37°C.

### Lipid peroxide detection in live cells

Lipid peroxides were detected using the cell-permeant dye Liperfluo (Dojindo Molecular Technologies) as described in (26). Cells grown in glass-bottom dishes (Ibidi) were loaded with 5 µM Liperfluo during the final 30 min of *t-*BOOH exposure and counterstained with NucBlue Live ReadyProbes reagent (Hoechst 33342). In some experiments, cells were pre-treated with the ferroportin inhibitor VIT-2763 (1 µM) for 2 h. Live-cell imaging was conducted using a Zeiss Axio Observer 7 inverted microscope fitted with a 20X/0.8 objective and Zen software (Zeiss). Fluorescence intensity was measured using FIJI v2.1.0 in at least 90 randomly selected cells across 9-15 fields in three replicates, averaged, and expressed as fold change relative to untreated controls for each species.

### Cell viability and cytotoxicity

Cytotoxicity and viability were evaluated via simultaneous use of GF-AFC and bis-AAF-R110 substrates (Promega). Cells were seeded overnight in fibronectin-coated clear-bottom 96-well plates at 10,000 cells per well and treated with increasing concentrations of *t-*BOOH for 2 h. Cells were then incubated for 30 min at 37°C with a cell-permeant substrate (GF-AFC), which fluoresces after cleavage by live-cell proteases, and a cell-impermeant substrate (bis-AAF-R110), which fluoresces after cleavage by proteases released from cells that have lost membrane integrity. Fluorescence was read at 400_Ex_/505_Em_ (viability) and 484_Ex_/520_Em_ (cytotoxicity) using a SpectraMax M3 plate reader (Molecular Devices). Data are expressed as fold change from untreated controls for each species. In separate experiments, cell viability and cytotoxicity were assessed using a commercial imaging kit (Thermo Fisher, catalog number: R37601), in cells treated with *t-*BOOH with and without ferrostatin-1 (10 µM), as previously described (26).

### Intracellular iron measurement

Cells seeded in glass-bottom dishes were rinsed three times with serum-free Live Cell Imaging Solution (Invitrogen), incubated with or without 1 µM VIT-2763 (Med Chem Express, cat. # HY-112220) for 2 h, and then treated with a solution containing 1 µM FerroOrange (Dojindo Molecular Technologies), with or without 300 µM *t*-BOOH, for 2 h. FerroOrange irreversibly fluoresces at 543_Ex_/580_Em_ upon reaction with ferrous iron. Cells were imaged using a Zeiss Axio Observer 7 inverted microscope equipped with a 20X/0.8 objective and Zen software (Zeiss). Fluorescence intensity was measured using FIJI v2.1.0 in 90 randomly selected cells across nine fields in three replicates, averaged, and expressed as fold change relative to untreated controls.

### BODIPY staining and lipid droplet quantification

Cells seeded in glass-bottom plates were treated with or without 100 µM *t-*BOOH for 2 h and stained with BODIPY 493/503 (ThermoFisher) diluted in PBS for 20 min, according to the manufacturer’s instructions. Live-cell imaging was performed using a Zeiss Axio Observer 7 inverted microscope system equipped with a 20X/0.8 objective and Zen software (Zeiss). Fluorescence intensity was quantified using FIJI v2.1.0 in at least five cells per field across 15 fields, with three replicates per condition. Fluorescence intensity in each field was normalized using a cell-free region of interest within that field. Results were expressed as fold change relative to untreated controls. The number of lipid droplets (LDs) in live cells was quantified using FIJI. Images were converted to 8-bit followed by contrast optimization. LDs were counted using the multi-point tool in at least three cells per field across 15 fields, with three replicates per species.

In some experiments, cells stained with BODIPY were fixed with 4% paraformaldehyde and imaged using a Zeiss LSM 880 FCS confocal microscope fitted with a 63X oil objective and Zen Black software (Zeiss). Images were imported into Huygens Professional Software (Scientific Volume Imaging, v26.04) for deconvolution to reduce image distortion using the Classic Maximum Likelihood Estimation (CMLE) algorithm with the theoretical point-spread function, a signal-to-noise ratio of 9.70, maximum iterations of 30, and a quality threshold of 0.01.

### Glutathione peroxidase activity

Total glutathione peroxidase (GPX) activity was measured in three replicates per species per condition using cumene hydroperoxide as a substrate with a glutathione peroxidase commercial kit (Cayman Chemical). GPX activity was normalized to protein concentration determined using a Rapid Gold BCA Protein Assay Kit (Pierce).

### Western blot

Cells were lysed in phosphate-buffered saline containing 1% Triton X-100 and 2% Halt Protease and Phosphatase Inhibitor Cocktail. Total protein concentration was determined using a BCA Rapid Gold Protein Assay Kit (Pierce). Proteins were diluted in lithium dodecyl sulfate buffer containing β-mercaptoethanol, heated at 70°C for 10 min, and resolved on SDS-PAGE gels before transfer to nitrocellulose membranes. Membranes were incubated overnight with antibodies against glutathione peroxidase 4 (GPX4; Abcam cat. # 125066; RRID:AB_10973901; 1:3,000), peroxiredoxin-6 (PRDX6; Cell Signaling Technology, cat. # 95336; RRID:AB_2800244; 1:1,000), 4-hydroxynonenal (4HNE, Abcam cat. # ab46545), or acyl-CoA synthetase long-chain family member 3 (ACSL3, Proteintech cat. # 20710-1-AP, 1:1,000). Proteins were visualized using IRDye 800CW secondary antibodies (LICOR) and a two-color near-infrared system (Azure 500, Azure Biosystems). Membranes were stripped and reprobed with an antibody against β-actin (Cell Signaling Technology, cat. # 4967; RRID:AB_330288; 1,1:000) or valosin-containing protein (VCP; Cell Signaling Technology, cat. # 2649; RRID:AB_2214629; 1:1,000). The intensity of individual bands was quantified using FIJI v2.1.0 and normalized to β-actin or VCP.

### Immunofluorescence

Immunofluorescence was performed as previously described (27, 28). Briefly, cells grown on glass-bottom plates were fixed, permeabilized, and incubated overnight with an anti-ferroportin antibody (FPN1, Proteintech, cat. # 26601-1-AP; RRID:AB_2880571; 1:500). Alexa Fluor Plus 488 secondary antibodies (ThermoFisher cat. # A32731) were used at 1:500. Nuclei were counterstained with Hoechst 33342. Cells were imaged using a Zeiss Axio Observer 7 microscope fitted with a 20X/0.8 objective and Zen software (Zeiss). Fluorescence intensity in 6-8 fields per well was quantified in three independent samples per condition and normalized to the number of cells per field. Data are expressed as fold change from the untreated control.

### Gene silencing

Seal-specific siRNAs were designed using the Life Technologies custom RNAi design tool. Cells were transfected with 25 pmol of siRNA targeting *ACSL3* or *SLC40A1* mRNAs, or with non-targeting control sequences (ThermoFisher, cat. # 4390843) using Lipofectamine RNAiMAX (ThermoFisher) according to the manufacturer’s instructions. Gene knockdown was confirmed 72 h post-transfection using RT-qPCR or western blot. siRNA sequences are listed in **Table S1**.

### RNA extraction and RT-qPCR

RNA was extracted using TRIzol (Invitrogen). Genomic DNA was removed using the TURBO DNA-free Kit (Ambion) according to the manufacturer’s instructions. cDNA was synthesized using a High-Capacity cDNA Reverse Transcription Kit (Applied Biosystems). RT-qPCR was conducted under the following conditions: 1 min at 95°C followed by 40 cycles of 10 or 20 sec at 95°C and 30 sec at 60°C. Primer sequences are listed in **Table S1**. Expression levels for target genes were normalized to the geometric mean of three housekeeping genes (*GAPDH, ACTB,* and *18S* or *GAPDH, ACTB,* and *ACTG1*). Relative mRNA expression (fold change) was calculated between treatment and control conditions using the 2^-ΔΔCt^ method.

### RNA-seq

RNA was extracted from cells exposed to normoxia (21% O_2_), hypoxia (1 h at 0.5% O_2_), hypoxia/reoxygenation (1 h at 0.5% O_2_ followed by 30 min normoxia), or *t-*BOOH (100 µM for 2 h) using an RNeasy kit (Qiagen) with on-column DNase treatment. gDNA removal was confirmed by the absence of genomic DNA amplification in RNA samples by PCR. RNA yield and RNA integrity number (RIN) were determined using an Agilent 2100 Bioanalyzer; all samples had a RIN > 9.1. Oligo(dT) bead enrichment of mRNA was followed by random fragmentation and cDNA synthesis using random hexamers. First-strand synthesis was followed by nick-translation generation of the second strand. Strands were then purified, terminal repaired, and poly(A)-tailed. After adapter ligation of poly(A)-captured mRNA, libraries underwent size selection and PCR enrichment. Three libraries were generated and sequenced for each condition across all species. Sequencing depth was 30M reads per sample on a NovaSeq platform (Illumina).

### Transcriptome analyses

The northern elephant seal (https://www.dnazoo.org/assemblies/Mirounga_angustirostris) genome was annotated as previously described (29). Reads were mapped to the elephant seal, human (GRCh38), or sheep (ARS-UI_Ramb_v2.0) genomes using STAR aligner v2.7.10a (30). Annotated gene counts were as follows: elephant seal, 14,088; human, 39,943; sheep, 14,528. A total of 11,830 genes were commonly annotated across all three species and used for subsequent comparative analyses. Transcript levels were quantified with RSEM v1.3.1 (31). For each species, genes differentially expressed (DE) between conditions were identified using EBSeq v1.34.0 under R v4.1.3 with a false discovery rate (FDR) < 0.05 (32). Gene set enrichment analyses (GSEA) were conducted on summary lists of genes DE in pairwise comparisons between conditions using the KEGG pathway database with FDR < 0.1 (33). Reactome Pathway enrichment analyses were conducted on summary lists of genes DE in pairwise comparisons between conditions using the Reactome Pathway database in Cytoscape v3.10.5 with FDR < 0.0001.

### Lipid extraction and LC-MS/MS

Cells were trypsinized, washed, resuspended in ice-cold PBS, transferred to glass collection tubes, and stored at −80°C until analysis. Lipid extraction and mass spectrometry were conducted at the UCLA Lipidomics Laboratory. Lipids were extracted using a modified Bligh and Dyer method (34). Prior to biphasic extraction, a standard mixture of 74 lipid standards (Avanti # 330820, 861809, 330729, 330727, 791642) was added to each sample. Following two successive extractions, the pooled organic layers were dried in a Thermo SpeedVac SPD300DDA using a ramp setting of 4°C to 35°C for 45 min, for a total run time of 90 min. Lipid samples were resuspended in 1:1 methanol/dichloromethane containing 10 mM ammonium acetate and transferred to robovials (Thermo # 10800107) for analysis. Samples were analyzed by direct infusion on a Sciex 5500 with a Differential Mobility Device (DMS) LC-MS/MS system using a targeted acquisition list comprising 1450 lipid species across 17 subclasses. The DMS was tuned with EquiSPLASH LIPIDOMIX (Avanti # 330731). Data analysis was performed using an in-house workflow. Instrument settings, MRM lists, and analysis methods were previously published in (35). Quantitative values were normalized to cell counts.

### Mass spectrometry data analysis

Downstream comparative lipidomic data analysis and visualization were conducted in R v4.2.3 in RStudio v2026.01.0392 using an integrated differential expression analysis pipeline for mass spectrometry data (36). Data were first filtered to include only lipid species detected in at least two replicates per group. To address the inherent heteroscedasticity of high-throughput data, variance-stabilizing normalization (vsn) was applied using the *vsn* package (37), while surrogate variable analysis (sva) was performed using the *sva* package (38) to account for latent technical noise and potential batch effects. Differential abundance was modeled using the *limma* framework (36, 39). *Limma* was selected for its ability to model the feature-wise mean-variance relationships inherent in mass spectrometry-based -omics data (36). Statistical significance was determined using the empirical Bayes (eBayes) method with a trended variance prior (40). *p-*values were adjusted for multiple comparisons using the Benjamini-Hochberg (BH)-based FDR. Lipids with a *p_adj* < 0.05 and absolute log_2_(fold change) ≥1 were considered differentially abundant.

To visualize the global lipidome profile in sheep and seal cells, a volcano plot was generated using the *EnhancedVolcano* package with lipids colored by their respective class. The top differentially abundant lipids were converted to normalized *z*-scores and visualized using the *pheatmap* package. Samples were organized using hierarchical clustering (Euclidean distance, complete linkage), and lipids were categorized by saturation level. Functional enrichment of lipids significantly upregulated in seal compared to sheep cells was conducted using LION/web to identify overrepresented lipid classes and biophysical properties (41). Enrichment results were visualized using the *ggplot2* package (42), ranking the top 20 terms by -log_10_(*p-*value). Differentially abundant lipids in *t*-BOOH-treated vs. untreated seal cells were determined using multiple *t*-tests with Holm-Šídák correction at a 5% FDR in GraphPad Prism v.11.0.0.

### Statistical analyses

Statistical analyses were performed using GraphPad Prism v.11.0.0. Normality was evaluated using D’Agostino-Pearson or Shapiro-Wilk tests, depending on sample size. All datasets met normality assumptions. Differences between groups with equal variance were assessed by independent two-tailed *t*-test or one-way ANOVA. ANOVA results were corrected for multiple comparisons using Šídák’s multiple comparisons test (α = 0.05). Multiple *t*-tests were corrected for multiple comparisons using the two-stage linear step-up procedure of Benjamini, Krieger and Yekutieli with a 5% FDR or Holm-Šídák method. Data are presented as mean ± SD unless otherwise indicated.

## Results

### Elephant seal cells evade lipid peroxidation

Elephant seals undergo repeated hypoxia-reoxygenation events during diving bouts, where arterial PO_2_ levels drop below 20 mmHg or near-complete oxygen depletion even during routine dives (7, 8). Hypoxia-reoxygenation increases oxidant generation, compromising tissue function in most species (43), but elephant seals are naturally adapted to cope with these extreme oxygen fluctuations (44). Therefore, we tested whether vascular endothelial cells derived from elephant seals can resist oxidative stress using a comparative framework. Treatment with hydroperoxide for 2 h increased cell death 2- to 3-fold in human and sheep but not seal cells **(Fig 1A)**, which also remained more viable after hydroperoxide treatment **(Fig S2)**. Previous work shows that seal tissues contain higher antioxidant levels than tissues from non-diving mammals (15, 45) and that hypoxia exposure stimulates GSH production in seal cells (24). Therefore, we considered whether GPX4 and peroxiredoxin 6 (PRDX6), two enzymes with reported phospholipid hydroperoxide glutathione peroxidase activity (14, 46) were higher in seal cells than in human cells. In contrast to previous observations at the tissue level (17) GPX4 and PRDX6 were 3-6× lower in seal cells than in human cells, while the basal abundance of 4-hydroxynonenal (4-HNE)-protein adducts was not different between species **(Fig. 1B)**. Similarly, total GPX activity was lower in seal than in human or sheep cells and was not induced upon exposure to 100 µM *t-*BOOH **(Fig 1C)**, suggesting that higher glutathione peroxidase activity is not the primary mechanism seals use to cope with hydroperoxide exposure. We then measured lipid peroxides using the Liperfluo probe (47) in live seal and human cells treated with hydroperoxide. Despite exhibiting low GPX4 and PRDX6, seal cells, but not human cells, remained protected against lipid peroxidation **(Fig 1D)**. Hydroperoxide treatment also failed to increase GPX4 or PRDX6 mRNA expression in seal cells **(Fig. 1E)**, whereas GPX4 protein levels remained unchanged in both species **(Fig. 1F)**. Consistent with previous reports, hydroperoxide treatment increased PRDX6 mRNA and protein levels in human cells (48) and this increase was also observed in seal cells at the protein level, although overall PRDX6 abundance remained lower in seal cells than in human cells **(Fig. 1F)**. These results suggest that while seal cells likely possess endogenous mechanisms to evade lipid peroxidation, these mechanisms might not include active lipid hydroperoxide scavenging.

**Figure 1.**
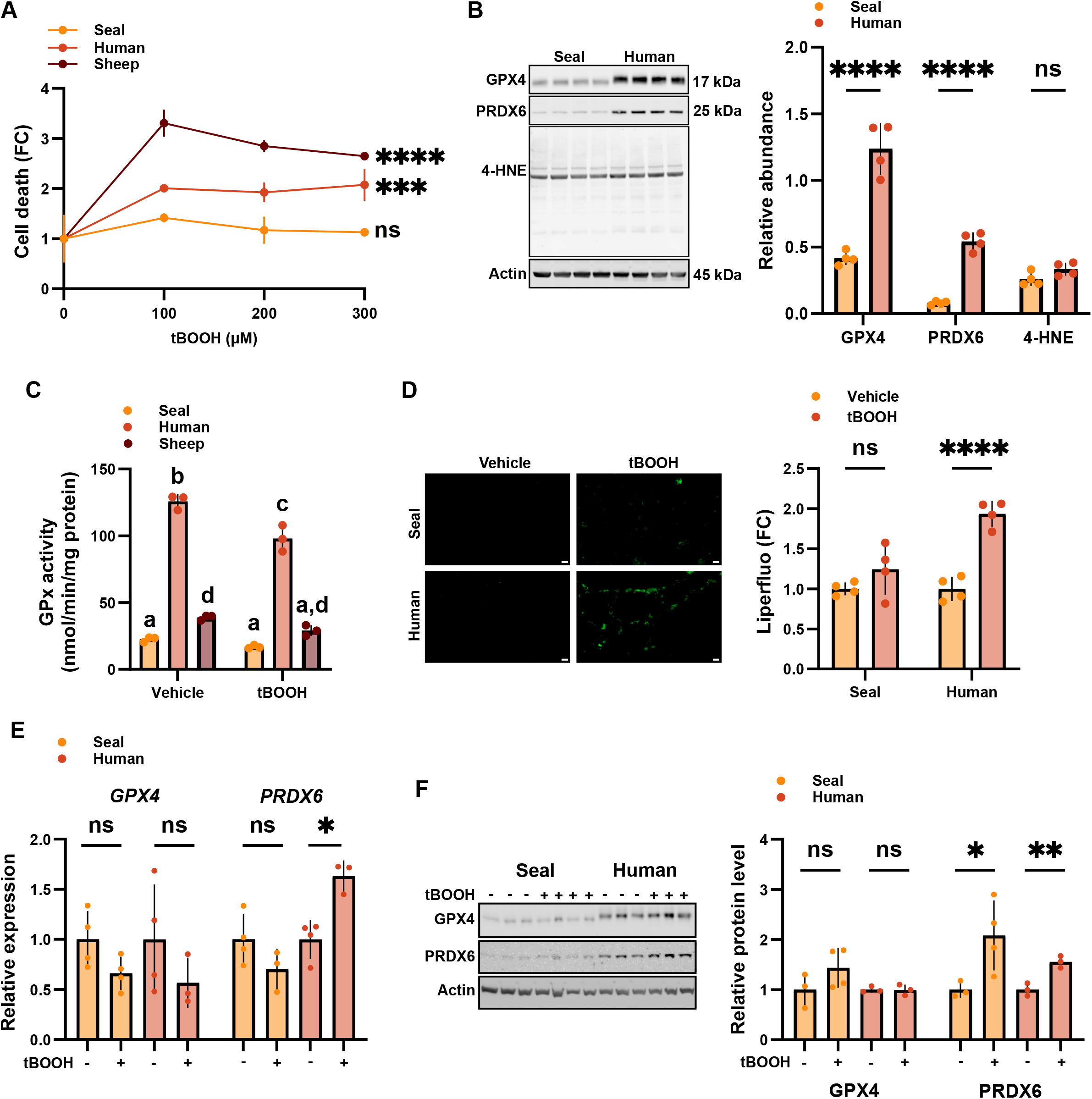
Hydroperoxide treatment does not induce lipid peroxidation in elephant seal cells. A) Fold change in cell death measured using bis-AAF-R110 in seal, human, and sheep endothelial cells treated with increasing concentrations of *t-*BOOH for 2 h. Data are mean ± SD, N = 3 per group. B) Western blot for GPX4, PRDX6, and 4-HNE. C) Total GPX activity measured using cumene hydroperoxide as a substrate in cells treated with 100 µM *t-*BOOH for 2 h. Different letters indicate statistical significance (p<0.05). D) Liperfluo fluorescence in cells treated with 300 µM *t-*BOOH for 1.5 h. E) GPX4 and PRDX6 mRNA and F) protein levels in cells treated with 100 µM *t-*BOOH for 2 h. * p<0.05, ** p<0.01, *** p<0.001, **** p<0.0001.

### Transcriptomic profiling links iron handling and lipid remodeling to oxidative stress resistance in seal cells

Our results suggest that active hydroperoxide scavenging is not a primary strategy seal cells use to evade lipid peroxidation. Therefore, we conducted a comparative transcriptomic analysis across seal, human, and sheep cells exposed to hypoxia, hypoxia-reoxygenation, or hydroperoxide to explore the mechanisms underlying lipid peroxidation resistance in seal cells. We found limited overlap in transcriptional programs across species **(Fig 2A)**, although condition-specific DE genes shared by all species generally displayed conserved expression patterns **(Fig S3)**. Hydroperoxide exposure upregulated gene expression in seal cells **(Fig S4)**, though the most extensive changes occurred with hypoxia exposure **(Fig 2B)**.

**Figure 2.**
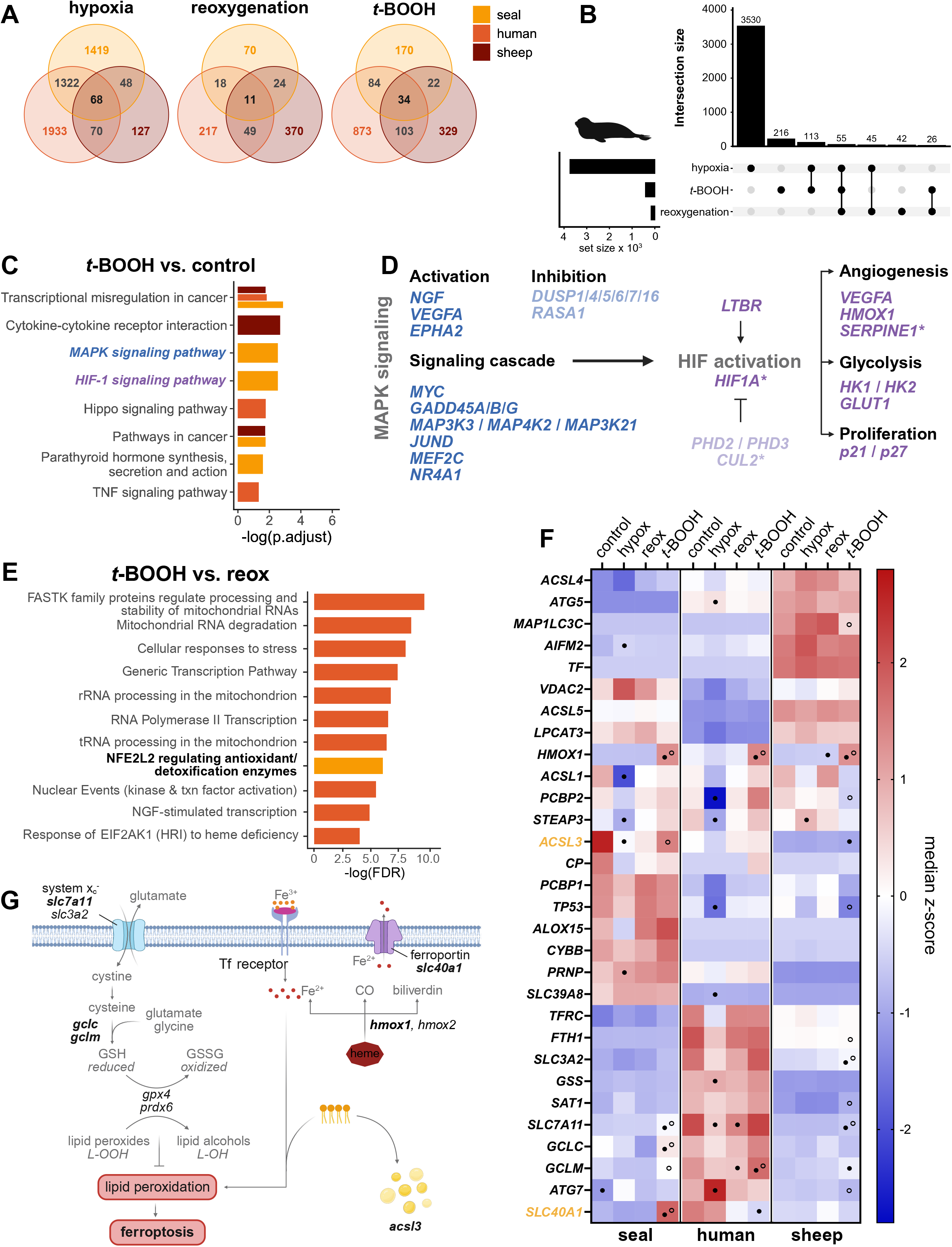
Transcriptional profiling links iron handling and lipid remodeling to lipid peroxidation resistance in seal cells. A) Number of genes DE vs. control in each species. Counts are constrained to genes identified as common among the annotations for all three species (11,830 total genes). B) UpSet plot showing the unique and common DEGs vs. control for seal. C) Selected KEGG enrichment results for genes DE in *t-*BOOH vs. control. D) Representation of MAPK activation of the HIF-1 pathway, including genes DE in seal cells treated with *t*-BOOH vs. control. Blue, MAPK signaling pathway; purple, HIF-1 signaling pathway. Lighter font colors indicate genes involved in pathway repression. * indicates downregulation. E) Reactome Pathway enrichments for genes DE in *t-*BOOH treatment compared to reoxygenation; FDR <0.0001. F) Expression of ferroptosis pathway genes present in seal enrichment in (E). Data are median *z*-scores for N = 3. Closed circles indicate genes DE in hypoxia, reoxygenation, or *t-*BOOH exposure vs. control. Open circles indicate genes DE in *t-*BOOH exposure vs. reoxygenation. G) Schematic representation of the ferroptosis pathway. Bold text indicates genes upregulated by *t-*BOOH exposure in seal cells from (F).

In seal cells, hydroperoxide exposure induced gene expression signatures associated with the MAPK and HIF-1 pathways **(Fig 2C; Fig S5)**, with all MAPK signaling genes upregulated **(Fig 2D)**. MAPK signaling activates HIF-1 via p38-mediated HIF-1α phosphorylation and p300 transactivation (49, 50). Among the HIF-1 pathway genes upregulated in seal cells, *HIF1A* mRNA expression was downregulated, and two prolyl hydroxylases (*PHD2, PHD3*) were upregulated (51). In contrast, *CUL2*, a scaffolding E3 ubiquitin ligase involved in HIF-1α degradation (52), was downregulated. Seal cells also upregulated *p21/p27*, which interacts with the pVHL subunit of the E3 ubiquitin ligase (53). Similarly, hydroperoxide treatment upregulated HIF-1 target genes involved in glycolysis, including two hexokinase isozymes, and *CD18*, which promotes leukocyte adhesion **(Fig 2D)**. Additionally, hydroperoxide exposure upregulated pro-angiogenic factors *VEGFA* and *HMOX1* while downregulating *SERPINE1,* which inhibits the proteolytic cascade that degrades the extracellular matrix during cell migration (54) **(Fig 2D)**. Together, these results suggest that hydroperoxide exposure induces a pseudo-hypoxic response in seal cells **(Fig S6)** (24).

In seal cells only, hydroperoxide exposure significantly enriched the NRF2 pathway **(Fig 2E)**. All NRF2 pathway-associated genes in this enrichment, including *NFE2L2*, *TXNRD1, SLC7A11, GSR, GCLC, and GCLM,* were upregulated by *t-*BOOH exposure, suggesting that antioxidant- and glutathione-mediated protection against oxidative stress may play a role despite limited evidence for GPX-mediated resistance under these conditions. Given that neither lipid peroxidation nor cell death increased with hydroperoxide treatment in seal cells **(Fig 1D)**, we also evaluated expression patterns in ferroptosis-associated genes (KEGG:hsa04216). Hydroperoxide exposure upregulated six hit genes in the pathway in seal cells when compared with hypoxia-reoxygenation **(Fig 2F)**. Of these six genes, three are involved in glutathione metabolism (*GCLC*, *GCLM, SLC7A11*), one is involved in lipid metabolism (*ACSL3*), and two are involved in iron homeostasis (*SLC40A1; HMOX1*) **(Fig 2G)**. *AIFM2* (*FSP1;* ferroptosis suppressor protein 1), an NRF2-mediated ferroptosis suppressor, was expressed at relatively low levels in seal cells. *SLC39A8 (ZIP8),* a divalent metal cation transporter involved in uptake of zinc, manganese, and iron, was highly expressed in seal cells compared with human and sheep but did not exhibit a response to any condition. *ACSL3* (acyl-CoA synthetase long-chain family member 3) expression was particularly high in seal cells compared to human cells or sheep cells, whereas only *SLC40A1* (*FPN1;* ferroportin-1) displayed a seal-specific expression pattern across treatments and was markedly upregulated in response to hydroperoxide exposure in seal cells only. These results are consistent with our data indicating that seal cells do not exhibit exceptionally high expression, abundance, or activity of lipid hydroperoxidases **(Fig 1B-F)**. Furthermore, these results suggest that seal endothelial cells may avoid lipid peroxidation by modulating iron metabolism and lipid composition.

### Lipid droplets shield seal cells from lipid peroxidation

Our comparative transcriptomic analyses showed higher basal *ACSL3* expression in seal cells than in human or sheep cells **(Fig 3A)**, which we confirmed by western blot **(Fig 3B)**. ACSL3 converts free long-chain fatty acids into fatty acyl-CoA esters, exerting cytoprotective effects by channeling monounsaturated fatty acids (MUFAs) to displace polyunsaturated fatty acids (PUFAs) from membranes and lower susceptibility to lipid peroxidation (55, 56). Accordingly, we conducted unbiased comparative lipidomic profiling in seal and sheep cells, which exhibit marked differences in ACSL3 levels and susceptibility to hydroperoxide treatment **(Fig 1A, 3A, 3B)**. Our results show that the seal cell lipidome is highly enriched in triglycerides (TGs) compared with sheep cells **(Fig 3C)**.

**Figure 3.**
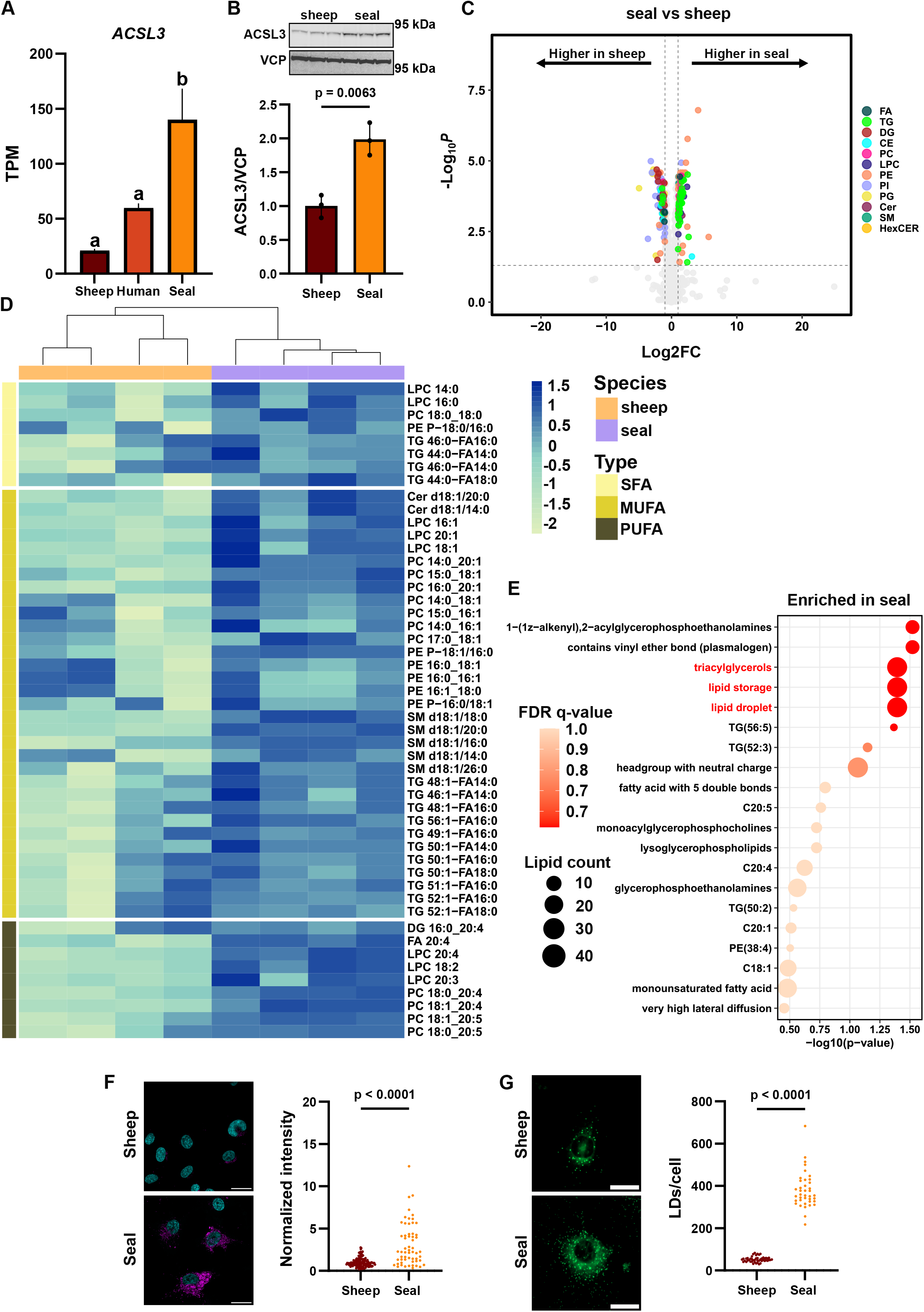
The elephant seal lipidome is enriched in triglycerides and MUFAs. A) *ACSL3* transcript levels in seal, human, and sheep cells. Data are mean ± SD (N = 3 per group). Different letters denote statistical significance. TPM, transcripts per million.B) Western blot analyses for ACSL3 in seal and sheep cells. C) Volcano plot showing differentially abundant lipids (in non-gray color, FC > 2, FDR < 0.05) in seal cells compared to sheep cells. FA, fatty acids; TG, triacylglycerol; DG, diacylglycerol; CE, cholesteryl ester; PC, phosphatidylcholine; LPC, lysophosphatidylcholine; PE, phosphatidylethanolamine; PI, phosphatidylinositol; PG, phosphatidylglycerol; Cer, ceramide; SM, sphingomyelin; HexCER, hexosylceramide. D) Heatmap of the top 50 differentially abundant lipids (normalized *Z*-scores) upregulated in elephant seal cells compared to sheep cells, showing hierarchical clustering of samples and lipids categorized by saturation level at 5% FDR. SFA: saturated fatty acids; MUFA: monounsaturated fatty acids; PUFA: polyunsaturated fatty acids. E) Lipid Ontology (LION/web) enrichment analysis in seal cells. F) BODIPY staining in sheep and seal cells. Cells were fixed and imaged using a confocal microscope. G) LD counts in live sheep and seal cells. Scale bars are 25 µm.

TG accumulation, high MUFA content, and ACSL3-mediated lipid droplet formation are strategies cells use to evade ferroptosis, a type of programmed cell death characterized by the uncontrolled accumulation of phospholipid hydroperoxides (55, 57, 58). Further examination of the lipid species enriched in seal cells showed a higher abundance of MUFA-containing phospholipids **(Fig 3D)**, whereas functional enrichment analysis showed the highest lipid counts and q-values for TGs, lipid stores, and lipid droplets **(Fig 3E)**, suggesting higher neutral lipid droplet content in seal cells than in sheep cells. Therefore, we quantified lipid droplet content in seal and sheep cells stained with BODIPY 493/503. BODIPY fluorescence intensity and lipid droplet count per cell were higher in seal cells than in sheep cells **(Fig 3F, 3G)**. Furthermore, mRNA levels of key enzymes that participate in TG biosynthesis and lipid droplet formation (*AGPAT1*, *FITM2*, and *GPAT3*) were consistently elevated in seal cells (59, 60) **(Fig S7)**.

Lipid droplets are crucial for storing neutral lipids, protecting cells against lipid peroxidation by sequestering easily oxidizable PUFAs (61). Hence, we investigated whether hydroperoxide treatment induces LD formation in seal cells. Using live-cell microscopy, we found that hydroperoxide treatment increases BODIPY fluorescence, suggesting the formation of neutral lipid droplets **(Fig 4A)**. Consistent with this observation, the expression of *PLIN3*, which participates in the formation and stabilization of nascent lipid droplets, and *GPAT4*, which catalyzes the initial, rate-limiting step of de novo TG synthesis in lipid droplets (62), also increased in response to hydroperoxide treatment **(Fig 4B)**. Similarly, hydroperoxide treatment increased the levels of cholesteryl esters (CEs) **(Fig 4C, 4D)**, neutral lipids stored within the core of lipid droplets, which can help buffer lipid peroxidation by sequestering PUFAs (61, 63). Consistent with this observation, PUFA-containing CEs (18:2 and 20:4) were the top two differentially abundant CEs in peroxide-treated seal cells, accompanied by increases across saturated and monounsaturated CEs (16:0, 16:1, 18:0, 18:1) **(Fig 4D)**.

**Figure 4.**
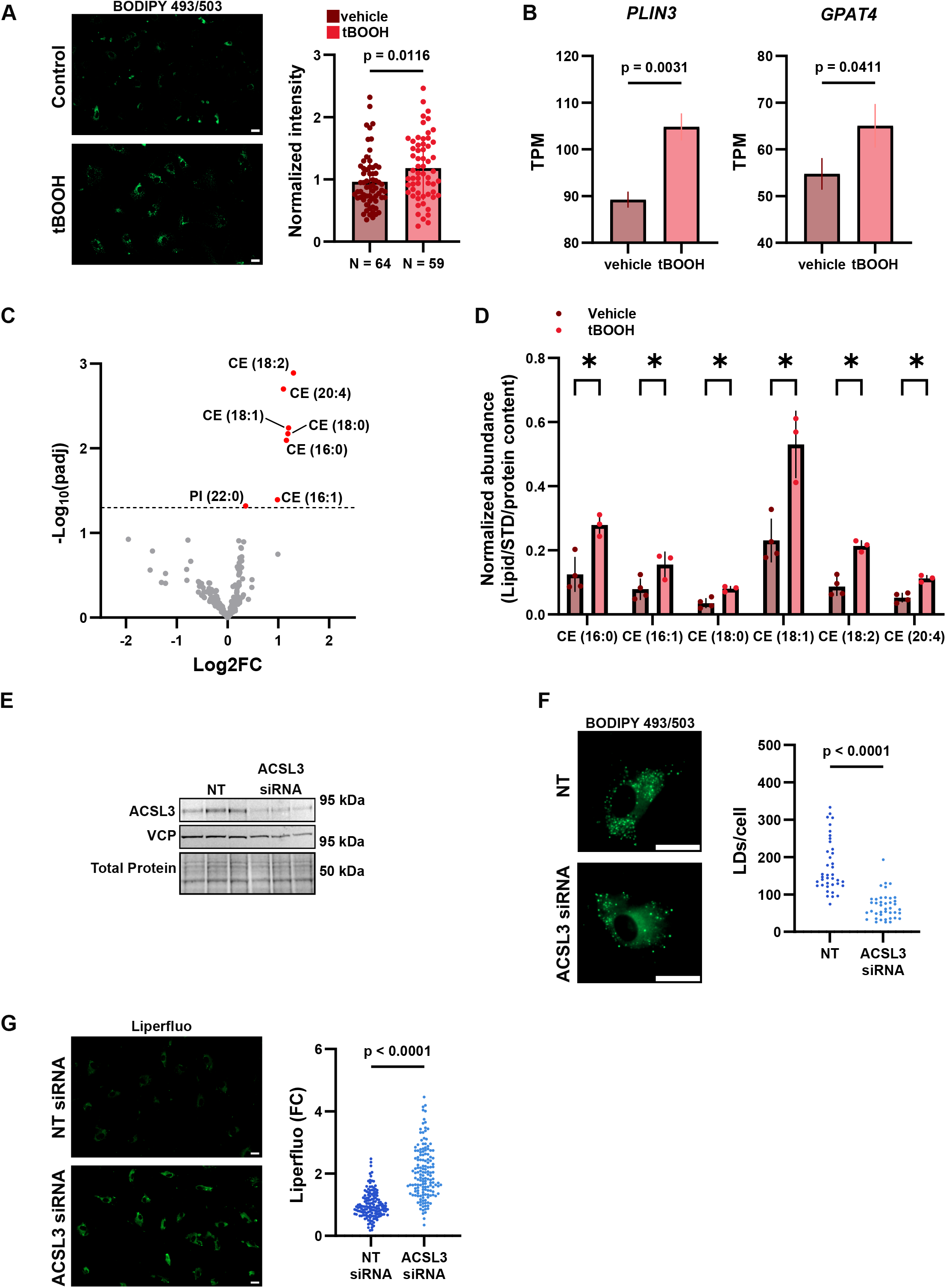
ACSL3-mediated lipid droplet formation protects elephant seal cells against lipid peroxidation. A) BODIPY staining in live seal cells treated with 100 µM *t*-BOOH for 2 h. B) *PLIN3* and *GPAT4* transcript levels in seal cells treated with *t-*BOOH. Data are mean ± SD (N = 3 per group). TPM, transcripts per million. C) Volcano plot of untargeted lipidomics data comparing seal cells treated with or without *t-*BOOH (FDR < 0.05). CE, cholesteryl esters. D) Cholesteryl ester differences in *t-*BOOH-treated vs untreated seal cells. Normalized raw data were compared using multiple t-tests with Holm-Šídák correction (* = p < 0.05). E) Western blot for ACSL3 in seal cells transfected with non-targeting (NT) or ACSL3 siRNAs. F) LD counts in ACSL3-depleted seal cells stained with BODIPY, N = 40 cells per group. G) Liperfluo fluorescence in ACSL3-depleted seal cells treated with 300 µM *t*-BOOH for 1.5 h, N = 150 cells per group. Scale bars are 25 µm.

ACSL3 promotes lipid droplet biogenesis by catalyzing the activation of long-chain fatty acids into acyl-CoAs, providing essential substrates for synthesizing and storing CEs within lipid droplets (64, 65). Therefore, we silenced ACSL3 to test whether this acyl-CoA synthetase drives lipid droplet formation in seal cells and whether ACSL3-depleted seal cells become susceptible to hydroperoxide-induced lipid peroxidation. ACSL3 depletion using siRNA **(Fig 4E)** decreased lipid droplet counts **(Fig 4F)** and sensitized seal cells to lipid peroxidation **(Fig 4G)**. Overall, these results show that ACSL3-driven lipid droplet accumulation contributes to seal cells’ resistance to lipid peroxidation.

### Seal cells actively export iron to evade lipid peroxidation

Our comparative transcriptomic analysis also showed that expression of ferroportin (*SLC40A1/FPN1*), the only known mammalian iron exporter (66), is highly responsive to hydroperoxide treatment in seal cells but not in human or sheep cells **(Fig 2F)**. Therefore, we evaluated FPN1 levels in hydroperoxide-treated seal cells by immunofluorescence. Hydroperoxide treatment increased FPN1 levels in seal cells **(Fig 5A)**, suggesting active iron export, which we confirmed using live-cell microscopy in cells loaded with FerroOrange, a fluorescent probe that enables intracellular iron measurement. Pre-treatment with VIT-2763, which promotes FPN1 ubiquitination, internalization, and degradation (67), reversed this effect **(Fig 5B)**.

**Figure 5.**
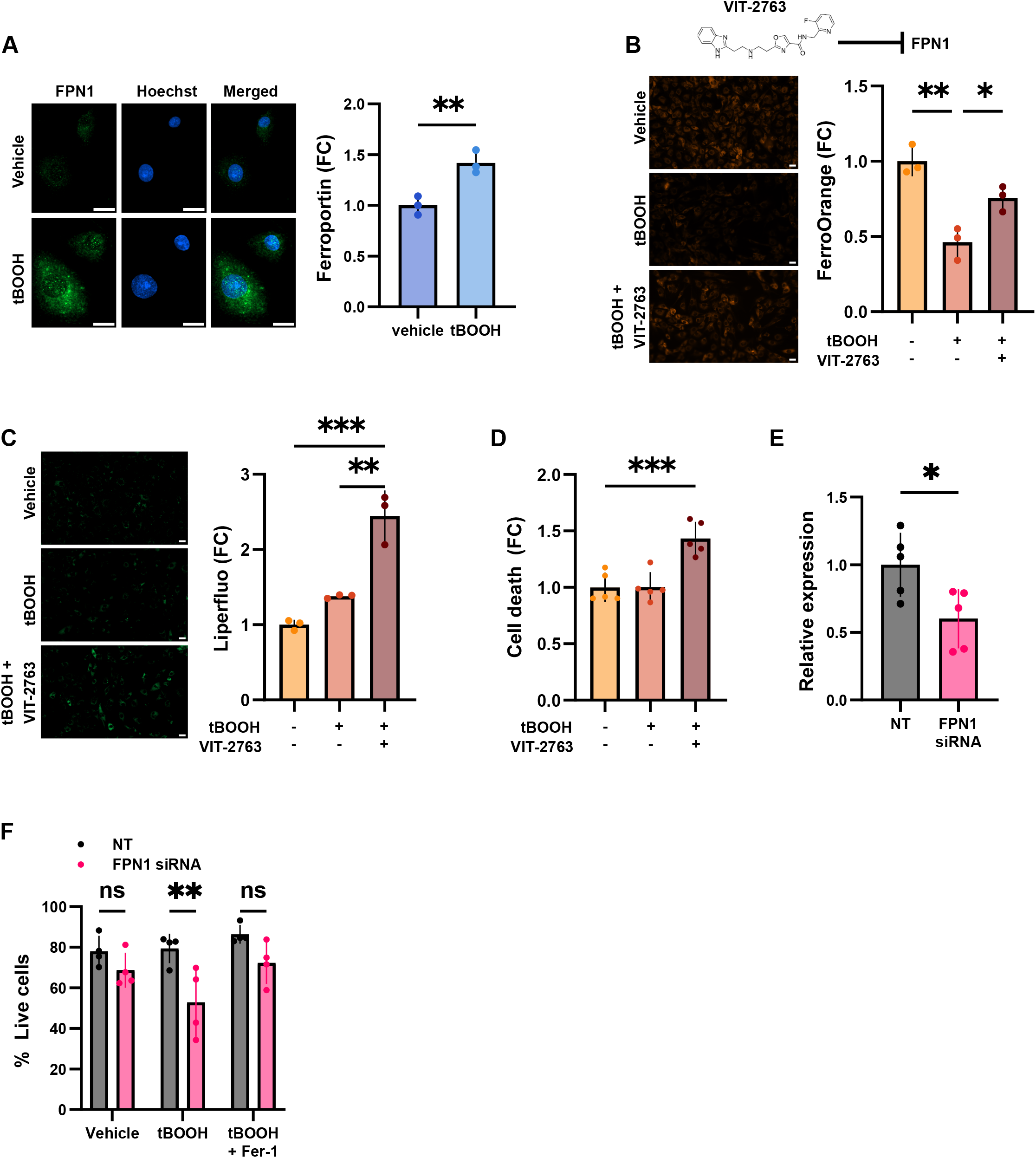
Elephant seal cells export intracellular iron to evade lipid peroxidation. A) Immunofluorescence analysis for FPN1 in seal cells treated with 100 µM *t*-BOOH for 2 h. B) Intracellular iron levels in cells loaded with FerroOrange (1 µM) and treated with 300 µM *t-*BOOH for 1.5 h, with or without the FPN1 inhibitor VIT-2763 (1 µM). C) Liperfluo fluorescence in cells treated with 300 µM *t*-BOOH for 1.5 h, with or without VIT-2763 (1 µM). D) Fold change in cell death measured using bis-AAF-R110 in seal cells treated with 300 µM *t-*BOOH for 1.5 h, with or without the FPN1 inhibitor VIT-2763 (1 µM). E) RT-qPCR for ferroportin in cells transfected with NT or FPN1 siRNAs. F) Cell viability/cytotoxicity in cells transfected with NT or ferroportin siRNAs and treated with 100 µM *t*-BOOH for 2 h with or without Ferrostatin-1 (10 µM). Data are mean ± SD. * p<0.05, ** p<0.01, *** p<0.001.

Intracellular iron drives lipid peroxidation by promoting the formation of reactive alkoxyl and peroxyl radicals (11). Thus, we tested whether blocking iron export by inhibiting FPN1 suppresses seal cell resistance to lipid peroxidation. Pre-treatment with VIT-2763 increased lipid peroxidation in hydroperoxide-treated seal cells **(Fig 5C)** to the levels observed in human cells **(Fig 1D)** and sensitized seal cells to hydroperoxide-induced cell death **(Fig 5D)**, further suggesting that exporting intracellular iron is a strategy seal cells use to evade lipid peroxidation. Consistent with this observation, FPN1 knockdown with siRNAs **(Fig 5E; 5F)** also sensitized seal cells to hydroperoxide-induced cell death (33% increase relative to non-targeting controls), whereas co-treatment with ferrostatin-1, a radical-trapping antioxidant that scavenges alkoxyl radicals (68), prevented this effect **(Fig 5E)**. These results show that, under oxidative stress, seal cells actively export iron via FPN1 to restrict the intracellular labile iron pool, thereby evading lipid peroxidation.

## Discussion

Elephant seals are consummate divers adapted to endure repetitive cycles of extreme hypoxia-reoxygenation. While previous studies show that elephant seals employ coordinated physiological adjustments to optimize oxygen use and upregulate antioxidant production under low-oxygen conditions (7, 8, 24, 44), the cellular mechanisms conferring oxidative stress tolerance to these extreme diving mammals remain incompletely understood. Here, we show that vascular endothelial cells derived from elephant seals are equipped to cope with hydroperoxide-induced lipid peroxidation. Multi-species transcriptomic profiling identified *ACSL3* and *FPN1* as central mediators of this protection. Further validation using comparative lipidomics alongside genetic and pharmacological approaches shows that elephant seal cells evade lipid peroxidation by forming lipid droplets and actively exporting intracellular iron.

Previous work shows that several antioxidant genes have undergone positive selection in diving mammals (16, 69, 70), whereas comparative studies between diving and non-diving species validate these genomic findings, revealing corresponding increases in antioxidant enzyme activities and protein levels (17, 18). We previously showed that seal cells, unlike human cells, efficiently conserve GSH under 1% oxygen, pointing to high basal antioxidant capacity as a critical defense against dive-associated oxidant generation (24). Here, however, we show that seal endothelial cells possess lower GPX activity and lower baseline GPX4 and PRDX6 levels. Therefore, our findings suggest that seal cells do not rely primarily on active lipid hydroperoxide scavenging to manage oxidative stress.

Our RNA-seq analysis revealed that hydroperoxide treatment induces a pseudo-hypoxic transcriptional signature in seal cells. Ischemic tolerance in seal tissues was first demonstrated over five decades ago (71), whereas subsequent work posited that seals limit oxidant generation by recycling purine metabolites accumulated during ischemia (72). More recent studies, however, suggest that oxidants generated during extended breath-holding may actually trigger an ischemic preconditioning response (44, 73, 74).

Our comparative transcriptomic analysis across seal, human, and sheep cells also showed that *ACLS3* expression is higher in seal than in human or sheep cells. ACSL3 suppresses lipid peroxidation by activating MUFAs and promoting their incorporation into membrane phospholipids (75), thereby displacing easily oxidizable PUFAs from cellular membranes. Additionally, ACSL3 promotes neutral lipid droplet biogenesis (55), which further shields cells from lipid peroxidation by sequestering PUFAs into stored TGs or cholesteryl esters, restricting their incorporation into membrane phospholipids and ultimately reducing overall membrane oxidizability (58, 59, 76). Consistent with these findings, comparative lipidomic profiling showed that seal cells are enriched in TGs and MUFAs. Furthermore, we found that seal cells have high baseline lipid droplet numbers, and hydroperoxide treatment induces additional neutral lipid formation while increasing the levels of CEs containing SFAs, MUFAs, and PUFAs, suggesting that PUFA sequestration in CE-enriched lipid droplets may protect seal cells against lipid peroxidation. ACSL3 depletion also reduced lipid droplet abundance and sensitized seal cells to hydroperoxide-induced lipid peroxidation, further suggesting that ACSL3-mediated lipid droplet formation contributes to seal cells’ tolerance to vascular oxidative stress.

Intracellular free iron catalyzes the Fenton and Fenton-like reactions, converting hydroperoxides into reactive radicals that drive lipid peroxidation propagation (11). Our results show that seal cells, but not human or sheep cells, upregulate the iron exporter FPN1 when exposed to hydroperoxide, and further live-cell imaging showed that seal cells actively export iron under these conditions. Furthermore, FPN1 inhibition promoted iron accumulation and lipid peroxidation, whereas FPN1 knockdown sensitized seal cells to hydroperoxide-induced cell death, further suggesting that under oxidant stress, seal cells export iron to evade lipid peroxidation. Consistent with our results, other work has shown that, in contrast to cancer cells, primary macrophages also upregulate FPN1 to prevent lipid peroxidation induced by incubation with RSL3 (77), which inhibits GPX4 (78, 79), TXNRD1 (80), and potentially other selenoproteins (81).

These findings should be interpreted in light of the traditional limitations of cell culture systems. Additionally, the upstream regulation of ferroportin induction remains unresolved but warrants further mechanistic investigation. Given that FPN1-mediated export redistributes rather than removes iron from the biological system, exported Fe^2+^ may be buffered *in vivo* through ferroxidase-dependent loading onto transferrin, uptake and storage in other tissues, and hepcidin-mediated feedback on FPN1 (66, 82). Although the degree to which these systems modulate iron flux in extreme breath-hold divers remains unknown, it is important to note that phocid seals exhibit remarkable physiological tolerance to high iron concentrations required for massive oxygen storage as well as high iron flux during lactation (83).

In summary, our results show that vascular endothelial cells derived from deep-diving elephant seals are naturally resistant to lipid peroxidation driven by high baseline concentrations of MUFAs and ACSL3-dependent lipid droplet content. Moreover, when exposed to hydroperoxides, seal cells dynamically respond by promoting further lipid formation and exporting iron via FPN1. Together, these coordinated strategies enable elephant seal cells to efficiently evade lipid peroxidation during acute oxidative stress.

## Acknowledgments

We thank Drs. Daniel Costa, Patrick Robinson, and Rachel Holser (University of California, Santa Cruz) for the collection and sharing of elephant seal placental tissue, Dr. Bret McNabb (UC Davis School of Veterinary Medicine) for providing sheep placental tissue, and Dr. Fung Lam for his assistance in obtaining human placental tissue. Dr. Kevin Williams, Baolong Su, and the UCLA Lipidomics core provided support for the lipidomics analysis. Drs. Denise Schichnes and Juliana Cho of the RCNR Biological Imaging Facility at UC Berkeley assisted with imaging training. Figures 2, S3, and S7 were created in BioRender and are licensed under agreement numbers JS2ACS675Z (Fig 2), SA2ACS67EX (Fig S3), UR2ACS67LK (Fig S4) and II2ACS67OF (graphical abstract) in accordance with CC BY 4.0. Figures can be viewed at https://BioRender.com/3bcslb7.

## Author Contributions

Conceived and designed research: KNA, ERP, JPV-M. Analyzed data: KNA, ERP, DDM-S, VEF, JPV-M. Performed experiments: KNA, ERP, DDM-S, ALL, DL, VEF, CDR, JPV-M. Interpreted results of experiments: KNA, ERP, JPV-M. Prepared figures: KNA, ERP, JPV-M. Drafted manuscript: KNA, ERP, JPV-M. Edited and revised manuscript: KNA, ERP, JPV-M. Approved final version: KNA, ERP, DDM-S, ALL, DL, VEF, CDR, JPV-M.

