## Supplementary material for "Elephant seal cells export iron and form lipid droplets to evade lipid peroxidation"

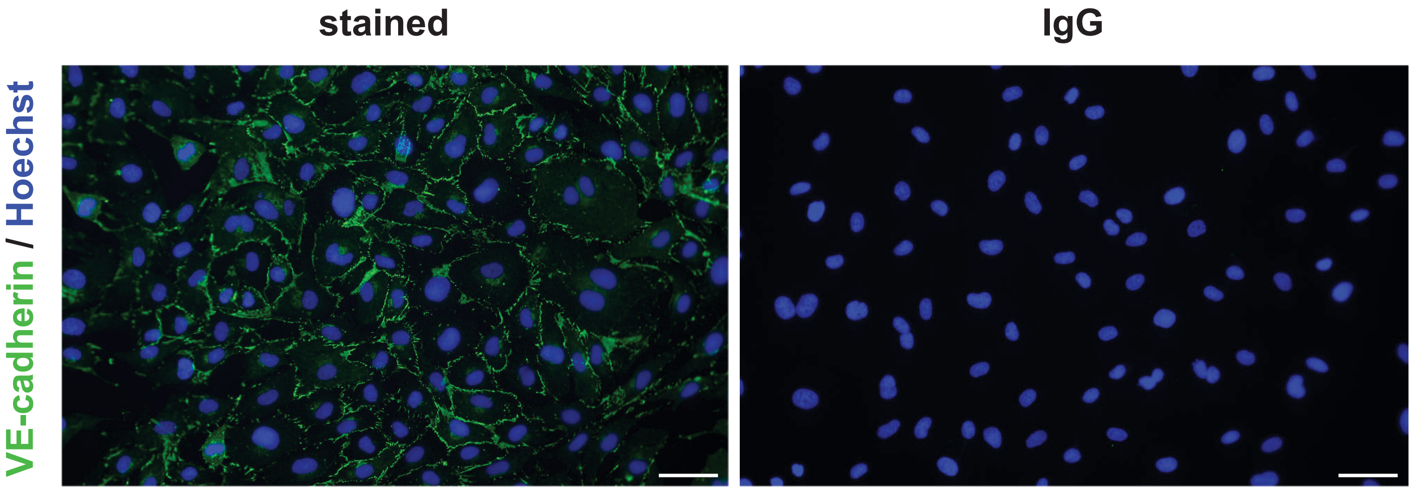
stained
IgG
VE-cadherin / Hoechst

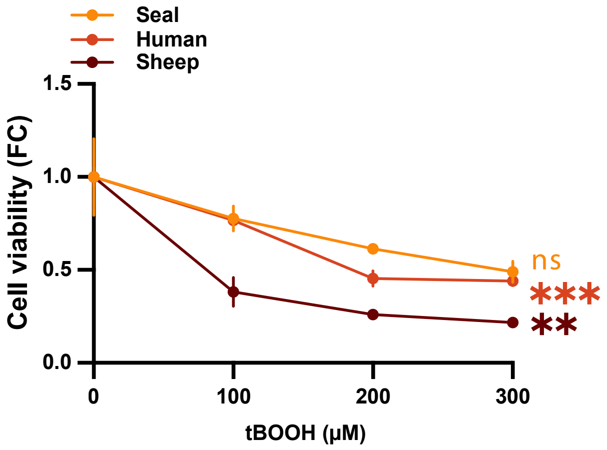

Seal
Human
Sheep
Cell viability (FC)
1.5
1.0
0.5
0.0
0
100
200
300
tBOOH (μM)
ns
***
**

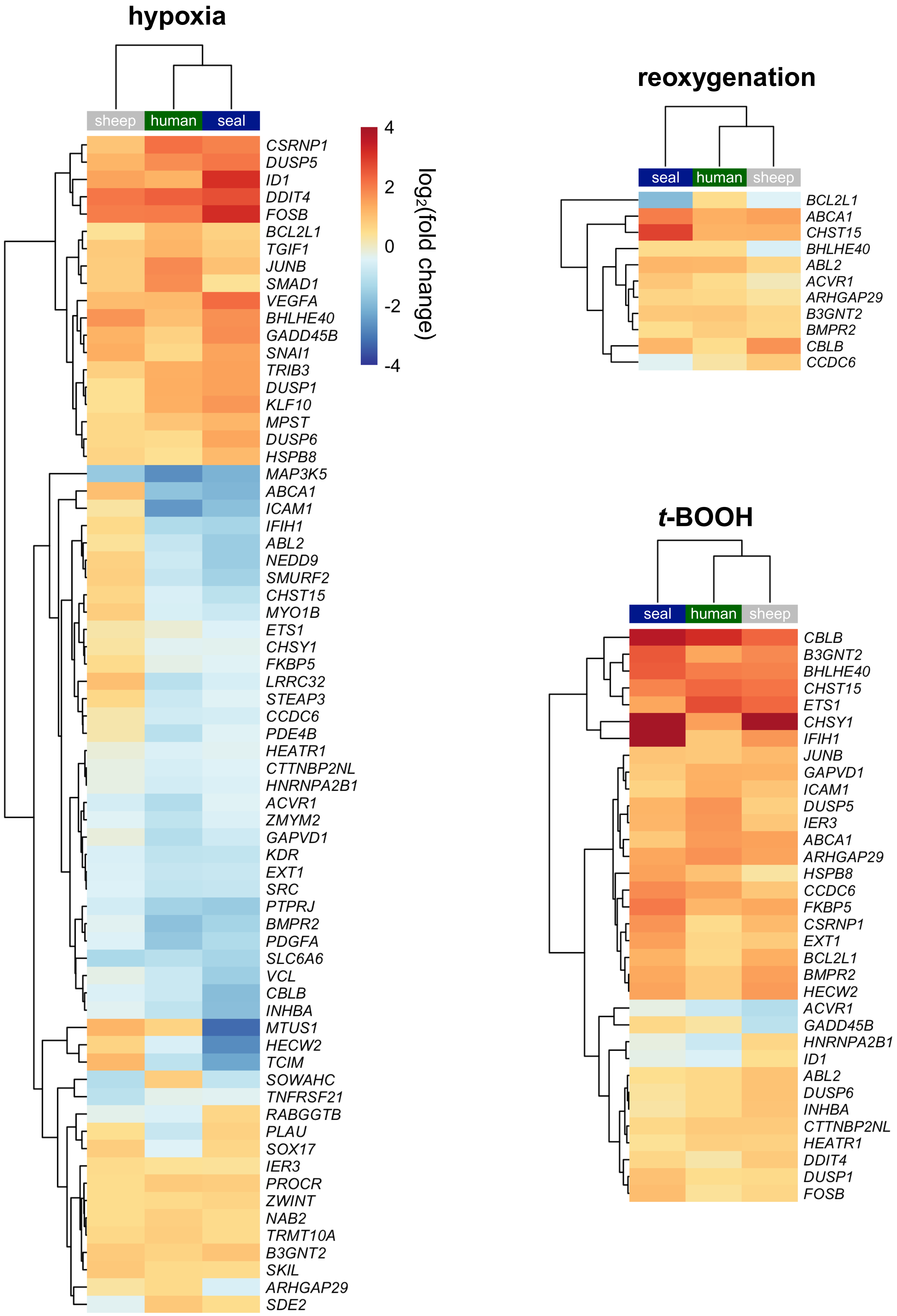
hypoxia
sheep
human
seal
CSRNP1
DUSP5
ID1
DDIT4
FOSB
BCL2L1
TGIF1
JUNB
SMAD1
VEGFA
BHLHE40
GADD45B
SNAI1
TRIB3
DUSP1
KLF10
MPST
DUSP6
HSPB8
MAP3K5
ABCA1
ICAM1
IFIH1
ABL2
NEDD9
SMURF2
CHST15
MYO1B
ETS1
CHSY1
FKBP5
LRRC32
STEAP3
CCDC6
PDE4B
HEATR1
CTTNBP2NL
HNRNPA2B1
ACVR1
ZMYM2
GAPVD1
KDR
EXT1
SRC
PTPRJ
BMPR2
PDGFA
SLC6A6
VCL
CBLB
INHBA
MTUS1
HECW2
TCIM
SOWAHC
TNFRSF21
RABGGTB
PLAU
SOX17
IER3
PROCR
ZWINT
NAB2
TRMT10A
B3GNT2
SKIL
ARHGAP29
SDE2
4
2
0
-2
-4
log2(fold change)
reoxygenation
seal
human
sheep
BCL2L1
ABCA1
CHST15
BHLHE40
ABL2
ACVR1
ARHGAP29
B3GNT2
BMPR2
CBLB
CCDC6
t-BOOH
seal
human
sheep
CBLB
B3GNT2
BHLHE40
CHST15
ETS1
CHSY1
IFIH1
JUNB
GAPVD1
ICAM1
DUSP5
IER3
ABCA1
ARHGAP29
HSPB8
CCDC6
FKBP5
CSRNP1
EXT1
BCL2L1
BMPR2
HECW2
ACVR1
GADD45B
HNRNPA2B1
ID1
ABL2
DUSP6
INHBA
CTTNBP2NL
HEATR1
DDIT4
DUSP1
FOSB

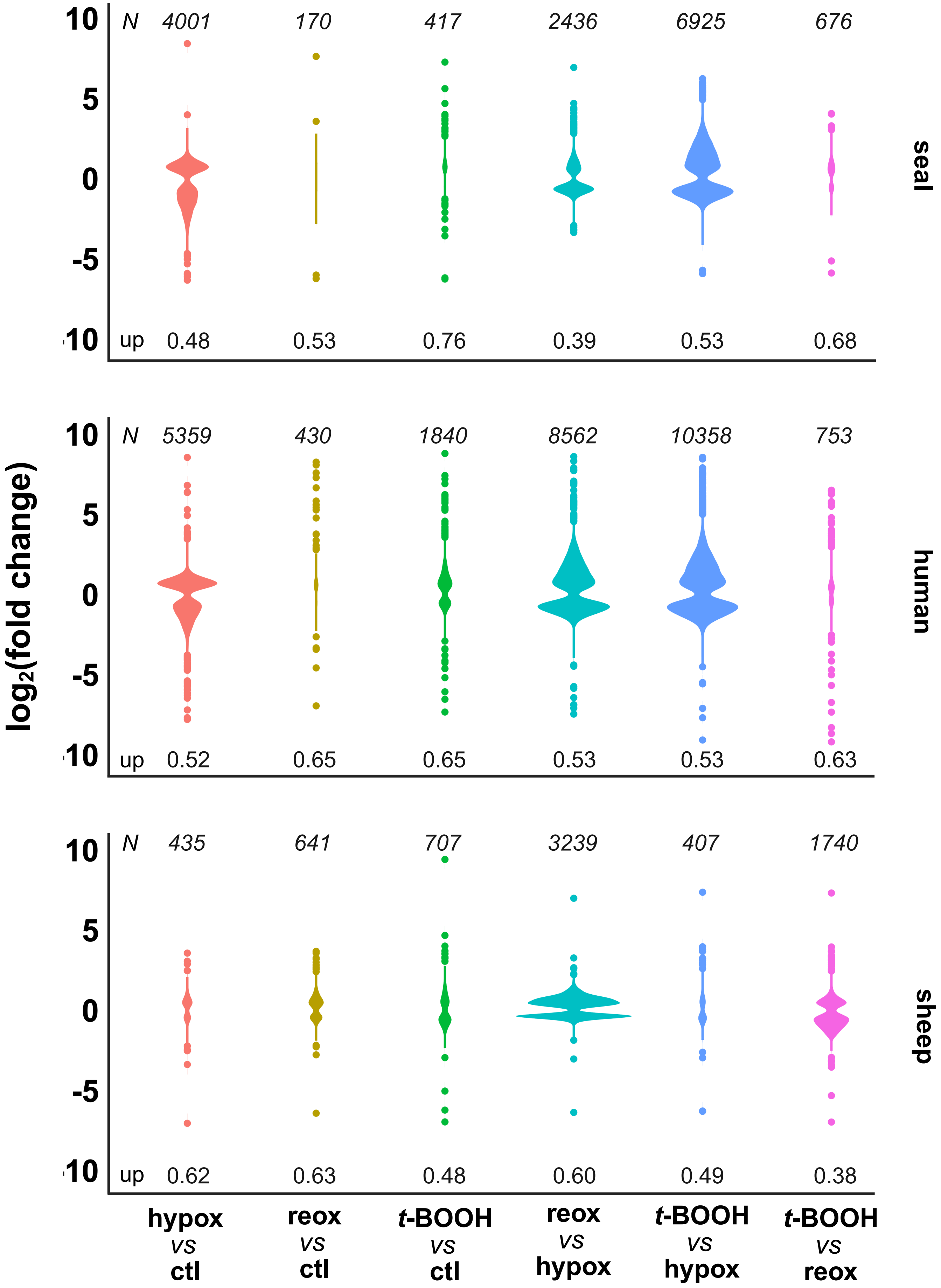

log2(fold change)
N 4001 170 417 2436 6925 676
up 0.48 0.53 0.76 0.39 0.53 0.68
seal
N 5359 430 1840 8562 10358 753
up 0.52 0.65 0.65 0.53 0.53 0.63
human
N 435 641 707 3239 407 1740
up 0.62 0.63 0.48 0.60 0.49 0.38
sheep
10
5
0
-5
-10
hypox vs ctl
reox vs ctl
t-BOOH vs ctl
reox vs hypox
t-BOOH vs hypox
t-BOOH vs reox

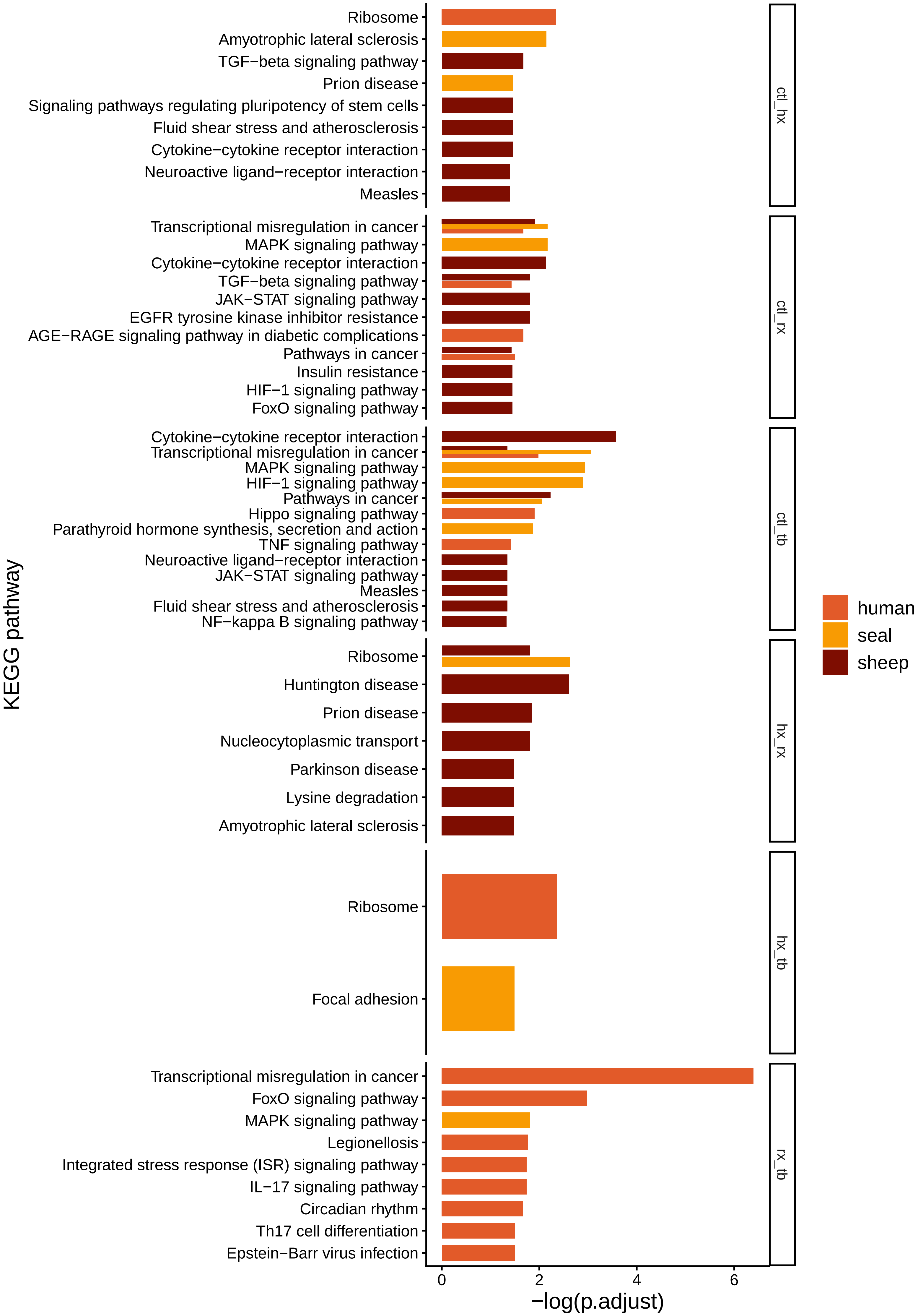
KEGG pathway
Ribosome
Amyotrophic lateral sclerosis
TGF−beta signaling pathway
Prion disease
Signaling pathways regulating pluripotency of stem cells
Fluid shear stress and atherosclerosis
Cytokine−cytokine receptor interaction
Neuroactive ligand−receptor interaction
Measles
ctl_hx
Transcriptional misregulation in cancer
MAPK signaling pathway
Cytokine−cytokine receptor interaction
TGF−beta signaling pathway
JAK−STAT signaling pathway
EGFR tyrosine kinase inhibitor resistance
AGE−RAGE signaling pathway in diabetic complications
Pathways in cancer
Insulin resistance
HIF−1 signaling pathway
FoxO signaling pathway
ctl_rx
Cytokine−cytokine receptor interaction
Transcriptional misregulation in cancer
MAPK signaling pathway
HIF−1 signaling pathway
Pathways in cancer
Hippo signaling pathway
Parathyroid hormone synthesis, secretion and action
TNF signaling pathway
Neuroactive ligand−receptor interaction
JAK−STAT signaling pathway
Measles
Fluid shear stress and atherosclerosis
NF−kappa B signaling pathway
ctl_tb
Ribosome
Huntington disease
Prion disease
Nucleocytoplasmic transport
Parkinson disease
Lysine degradation
Amyotrophic lateral sclerosis
hx_rx
Ribosome
Focal adhesion
hx_tb
Transcriptional misregulation in cancer
FoxO signaling pathway
MAPK signaling pathway
Legionellosis
Integrated stress response (ISR) signaling pathway
IL−17 signaling pathway
Circadian rhythm
Th17 cell differentiation
Epstein−Barr virus infection
rx_tb
0
2
4
6
−log(p.adjust)
human
seal
sheep

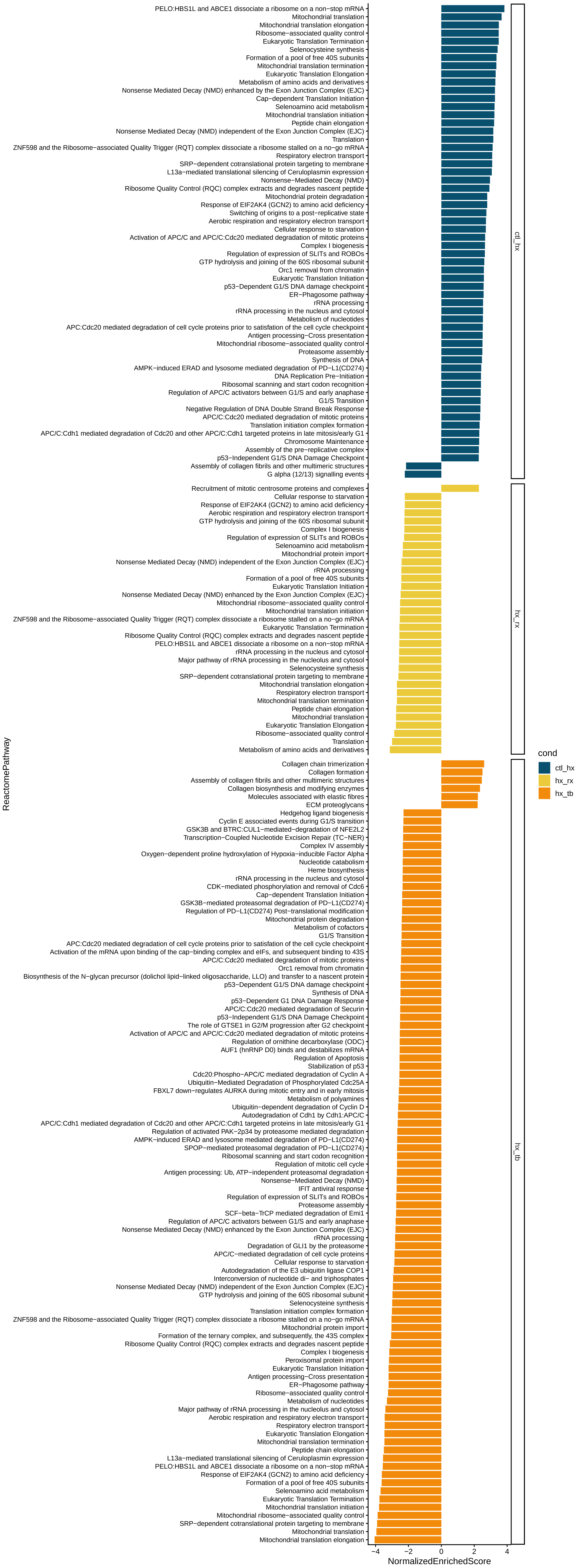
PELO:HBS1L and ABCE1 dissociate a ribosome on a non−stop mRNA
Mitochondrial translation
Mitochondrial translation elongation
Ribosome−associated quality control
Eukaryotic Translation Termination
Selenocysteine synthesis
Formation of a pool of free 40S subunits
Mitochondrial translation termination
Eukaryotic Translation Elongation
Metabolism of amino acids and derivatives
Nonsense Mediated Decay (NMD) enhanced by the Exon Junction Complex (EJC)
Cap−dependent Translation Initiation
Selenoamino acid metabolism
Mitochondrial translation initiation
Peptide chain elongation
Nonsense Mediated Decay (NMD) independent of the Exon Junction Complex (EJC)
Translation
ZNF598 and the Ribosome−associated Quality Trigger (RQT) complex dissociate a ribosome stalled on a no−go mRNA
Respiratory electron transport
SRP−dependent cotranslational protein targeting to membrane
L13a−mediated translational silencing of Ceruloplasmin expression
Nonsense−Mediated Decay (NMD)
Ribosome Quality Control (RQC) complex extracts and degrades nascent peptide
Mitochondrial protein degradation
Response of EIF2AK4 (GCN2) to amino acid deficiency
Switching of origins to a post−replicative state
Aerobic respiration and respiratory electron transport
Cellular response to starvation
Activation of APC/C and APC/C:Cdc20 mediated degradation of mitotic proteins
Complex I biogenesis
Regulation of expression of SLITs and ROBOs
GTP hydrolysis and joining of the 60S ribosomal subunit
Orc1 removal from chromatin
Eukaryotic Translation Initiation
p53−Dependent G1/S DNA damage checkpoint
ER−Phagosome pathway
rRNA processing
rRNA processing in the nucleus and cytosol
Metabolism of nucleotides
APC:Cdc20 mediated degradation of cell cycle proteins prior to satisfaction of the cell cycle checkpoint
Antigen processing−Cross presentation
Mitochondrial ribosome−associated quality control
Proteasome assembly
Synthesis of DNA
AMPK−induced ERAD and lysosome mediated degradation of PD−L1(CD274)
DNA Replication Pre−Initiation
Ribosomal scanning and start codon recognition
Regulation of APC/C activators between G1/S and early anaphase
G1/S Transition
Negative Regulation of DNA Double Strand Break Response
APC/C:Cdc20 mediated degradation of mitotic proteins
Translation initiation complex formation
APC/C:Cdh1 mediated degradation of Cdc20 and other APC/C:Cdh1 targeted proteins in late mitosis/early G1
Chromosome Maintenance
Assembly of the pre−replicative complex
p53−Independent G1/S DNA Damage Checkpoint
Assembly of collagen fibrils and other multimeric structures
G alpha (12/13) signalling events
ctl_hx
Recruitment of mitotic centrosome proteins and complexes
Cellular response to starvation
Response of EIF2AK4 (GCN2) to amino acid deficiency
Aerobic respiration and respiratory electron transport
GTP hydrolysis and joining of the 60S ribosomal subunit
Complex I biogenesis
Regulation of expression of SLITs and ROBOs
Selenoamino acid metabolism
Mitochondrial protein import
Nonsense Mediated Decay (NMD) independent of the Exon Junction Complex (EJC)
rRNA processing
Formation of a pool of free 40S subunits
Eukaryotic Translation Initiation
Nonsense Mediated Decay (NMD) enhanced by the Exon Junction Complex (EJC)
Mitochondrial ribosome−associated quality control
Mitochondrial translation initiation
ZNF598 and the Ribosome−associated Quality Trigger (RQT) complex dissociate a ribosome stalled on a no−go mRNA
Eukaryotic Translation Termination
Ribosome Quality Control (RQC) complex extracts and degrades nascent peptide
PELO:HBS1L and ABCE1 dissociate a ribosome on a non−stop mRNA
rRNA processing in the nucleus and cytosol
Major pathway of rRNA processing in the nucleolus and cytosol
Selenocysteine synthesis
SRP−dependent cotranslational protein targeting to membrane
Mitochondrial translation elongation
Respiratory electron transport
Mitochondrial translation termination
Peptide chain elongation
Mitochondrial translation
Eukaryotic Translation Elongation
Ribosome−associated quality control
Translation
Metabolism of amino acids and derivatives
hx_rx
Collagen chain trimerization
Collagen formation
Assembly of collagen fibrils and other multimeric structures
Collagen biosynthesis and modifying enzymes
Molecules associated with elastic fibres
ECM proteoglycans
Hedgehog ligand biogenesis
Cyclin E associated events during G1/S transition
GSK3B and BTRC:CUL1−mediated−degradation of NFE2L2
Transcription−Coupled Nucleotide Excision Repair (TC−NER)
Complex IV assembly
Oxygen−dependent proline hydroxylation of Hypoxia−inducible Factor Alpha
Nucleotide catabolism
Heme biosynthesis
rRNA processing in the nucleus and cytosol
CDK−mediated phosphorylation and removal of Cdc6
Cap−dependent Translation Initiation
GSK3B−mediated proteasomal degradation of PD−L1(CD274)
Regulation of PD−L1(CD274) Post−translational modification
Mitochondrial protein degradation
Metabolism of cofactors
G1/S Transition
APC:Cdc20 mediated degradation of cell cycle proteins prior to satisfaction of the cell cycle checkpoint
Activation of the mRNA upon binding of the cap−binding complex and eIFs, and subsequent binding to 43S
APC/C:Cdc20 mediated degradation of mitotic proteins
Orc1 removal from chromatin
Biosynthesis of the N−glycan precursor (dolichol lipid−linked oligosaccharide, LLO) and transfer to a nascent protein
p53−Dependent G1/S DNA damage checkpoint
Synthesis of DNA
p53−Dependent G1 DNA Damage Response
APC/C:Cdc20 mediated degradation of Securin
p53−Independent G1/S DNA Damage Checkpoint
The role of GTSE1 in G2/M progression after G2 checkpoint
Activation of APC/C and APC/C:Cdc20 mediated degradation of mitotic proteins
Regulation of ornithine decarboxylase (ODC)
AUF1 (hnRNP D0) binds and destabilizes mRNA
Regulation of Apoptosis
Stabilization of p53
Cdc20:Phospho−APC/C mediated degradation of Cyclin A
Ubiquitin−Mediated Degradation of Phosphorylated Cdc25A
FBXL7 down−regulates AURKA during mitotic entry and in early mitosis
Metabolism of polyamines
Ubiquitin−dependent degradation of Cyclin D
Autodegradation of Cdh1 by Cdh1:APC/C
APC/C:Cdh1 mediated degradation of Cdc20 and other APC/C:Cdh1 targeted proteins in late mitosis/early G1
Regulation of activated PAK−2p34 by proteasome mediated degradation
AMPK−induced ERAD and lysosome mediated degradation of PD−L1(CD274)
SPOP−mediated proteasomal degradation of PD−L1(CD274)
Ribosomal scanning and start codon recognition
Regulation of mitotic cell cycle
Antigen processing: Ub, ATP−independent proteasomal degradation
Nonsense−Mediated Decay (NMD)
IFIT antiviral response
Regulation of expression of SLITs and ROBOs
Proteasome assembly
SCF−beta−TrCP mediated degradation of Emi1
Regulation of APC/C activators between G1/S and early anaphase
Nonsense Mediated Decay (NMD) enhanced by the Exon Junction Complex (EJC)
rRNA processing
Degradation of GLI1 by the proteasome
APC/C−mediated degradation of cell cycle proteins
Cellular response to starvation
Autodegradation of the E3 ubiquitin ligase COP1
Interconversion of nucleotide di− and triphosphates
Nonsense Mediated Decay (NMD) independent of the Exon Junction Complex (EJC)
GTP hydrolysis and joining of the 60S ribosomal subunit
Selenocysteine synthesis
Translation initiation complex formation
ZNF598 and the Ribosome−associated Quality Trigger (RQT) complex dissociate a ribosome stalled on a no−go mRNA
Mitochondrial protein import
Formation of the ternary complex, and subsequently, the 43S complex
Ribosome Quality Control (RQC) complex extracts and degrades nascent peptide
Complex I biogenesis
Peroxisomal protein import
Eukaryotic Translation Initiation
Antigen processing−Cross presentation
ER−Phagosome pathway
Ribosome−associated quality control
Metabolism of nucleotides
Major pathway of rRNA processing in the nucleolus and cytosol
Aerobic respiration and respiratory electron transport
Respiratory electron transport
Eukaryotic Translation Elongation
Mitochondrial translation termination
Peptide chain elongation
L13a−mediated translational silencing of Ceruloplasmin expression
PELO:HBS1L and ABCE1 dissociate a ribosome on a non−stop mRNA
Response of EIF2AK4 (GCN2) to amino acid deficiency
Formation of a pool of free 40S subunits
Selenoamino acid metabolism
Eukaryotic Translation Termination
Mitochondrial translation initiation
Mitochondrial ribosome−associated quality control
SRP−dependent cotranslational protein targeting to membrane
Mitochondrial translation
Mitochondrial translation elongation
hx_tb
ReactomePathway
−4
−2
0
2
4
NormalizedEnrichedScore
cond
ctl_hx
hx_rx
hx_tb

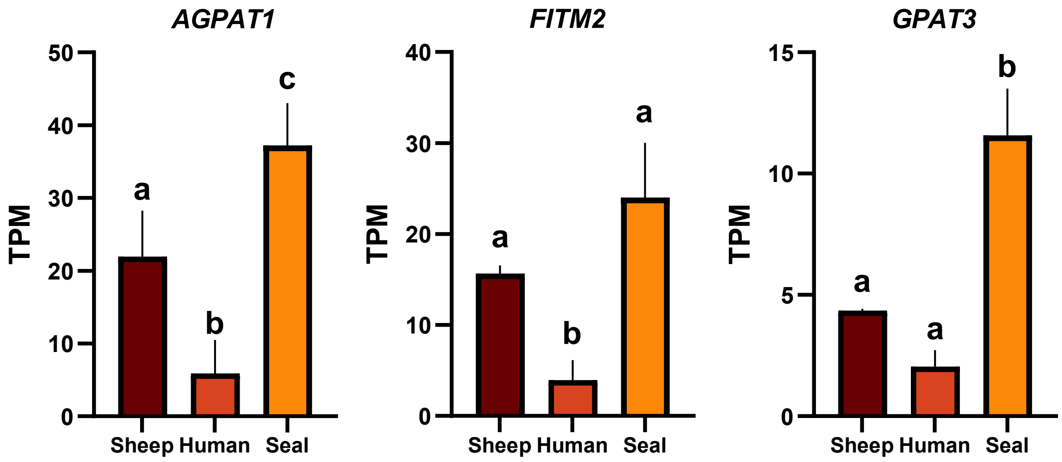
AGPAT1
TPM
50
40
30
20
10
0
a
b
c
Sheep
Human
Seal
FITM2
TPM
40
30
20
10
0
a
b
a
Sheep
Human
Seal
GPAT3
TPM
15
10
5
0
a
a
b
Sheep
Human
Seal

**SUPPLEMENTARY TABLES**

**Table S1. Nucleotide sequences**

| **siRNA** | | |
|---|---|---|
| **species** | **target gene** | **sequence 5' to 3' (sense / antisense)** |
| **Northern elephant seal *(Mirounga angustirostris)*** | ***ACSL3*** | AGGUUAUUCUUGGAAACUAtt |
| | | UAGUUUCCAAGAAUAACCUtt |
| | ***SLC40A1*** | AGAUGUAUUCGUUUGAUUUtt |
| | | AAAUCAAACGAAUACAUCUgt |
| **Primers** | | |
| **species** | **target gene** | **sequence 5' to 3' (forward / reverse)** |
| **Northern elephant seal *(Mirounga angustirostris)*** | ***GAPDH*** | CAA GGC TGA GAA CGG GAA GC |
| | | ATC GGC AGA GGG AGC AGA GA |
| | ***β-ACTIN*** | GAC ATC CGC AAG GAC CTC TA |
| | | GCC TCC AAT CCA CAC AGA GT |
| | ***γ-ACTIN*** | CGG TCA GTT CGT GGC TGA GG |
| | | AAG GCT CGG ACC TTC CCA AC |
| | ***GPX4*** | AAG TAC CGG GGC TTC GTG TG |
| | | CCA GCG GCG AAC TCT TTG AT |
| | ***PRDX6*** | GCA CCA CAG AGC TTG GCA GA |
| | | AGG ATG GCA AGG TCC CGA TT |
| | ***SLC40A1*** | CGG TCA TCC TGT GTG GGA TC |
| | | GTC TCC TCC TGC AAC GAC AA |
| **Human *(Homo sapiens)*** | ***GAPDH*** | CCT GCA CCA CCA ACT GCT TA |
| | | GGC CAT CCA CAG TCT TCT GAG |
| | ***β-ACTIN*** | TGA TGG TGG GCA TGG GTC AGA A |
| | | TCT GGG TCA TCT TCT CGC GGT T |
| | ***GPX4*** | AGA GAT CAA AGA GTT CGC CGC |
| | | TCT TCA TCC ACT TCC ACA GCG |
| | ***PRDX6*** | CGT GTG GTG TTT GTT TTT GG |
| | | CCA TCA CAC TAT CCC CAT CC |
| **All** | ***18S rRNA*** | GTA ACC CGT TGA ACC CCA TT |
| | | CCA TCC AAT CGG TAG TAG CG |

**Table S2. NFE2L2-regulated antioxidant/detoxification genes differentially expressed in seal cells during *t*-BOOH exposure compared with hypoxia-reoxygenation.**

| ***Gene name*** | **$log_2$(fold change)** |
|---|---|
| *HMOX1* | 4.04 |
| *TXNRD1* | 0.72 |
| *NQO1* | 0.91 |
| *SLC7A11* | 1.26 |
| *GSR* | 0.53 |
| *GCLC* | 1.06 |
| *NFE2L2* | 0.43 |
| *GCLM* | 0.87 |
| *BACH1* | 0.34 |
| *CREBBP* | 1.03 |

**SUPPLEMENTARY FIGURE LEGENDS**

**Figure S1. Primary arterial endothelial cells derived from sheep express the canonical endothelial marker VE-cadherin (CD144).** Green, VE-cadherin; blue, Hoechst. Scale bar is 50 µm.

**Figure S2. Peroxide treatment induces mild declines in viability in seal endothelial cells.** Fold change in viability measured using GF-AFC in seal, human, and sheep endothelial cells treated with increasing concentrations of *t*-BOOH for 2 h. Data are mean ± SD, *n*=3. Two-way ANOVA. ** p<0.01, *** p<0.001.

**Figure S3. DE genes shared between species for each condition display similar expression patterns.** Fold change data are calculated vs. control for each species x condition. Heatmaps of $log_2$(fold change) for were generated using the *pheatmap* package v1.0.13 in R v4.4.1. Genes and species were hierarchically clustered using Euclidean distance and complete linkage, without additional scaling. Gene sets are drawn from center segments of Venn diagrams in **Fig 2A**.

**Figure S4. Global gene expression patterns during hypoxia, hypoxia/reoxygenation, and *t*-BOOH treatment.** Italicized *N* above each plot indicates the total number of genes DE in a given comparison. Values below each plot indicate the proportion of DE genes upregulated in a given comparison.

**Figure S5. Complete KEGG pathway enrichment results for DE genes.** Gold, seal; orange, human; dark red, sheep. ctl, control; hx, hypoxia; rx, hypoxia-reoxygenation; tb, *t*-BOOH.

**Figure S6. Complete GSEA results for genes DE in seal cells.** Normalized Enrichment Score was calculated using Cytoscape and indicates net impact on pathway function (NES > 0, upregulated; NES < 0, downregulated). FDR < 0.001 for all pathways. No pathways were enriched for ctl_rx, ctl_tb or rx_tb. ctl, control; hx, hypoxia; rx, hypoxia-reoxygenation; tb, *t*-BOOH.

**Figure S7.** Baseline transcript levels of *AGPAT1*, *FITM2*, and *GPAT3* in seal, human, and sheep cells. Data are mean ± SD, *n*=3. Different letters denote statistical significance (p<0.05). TPM, transcripts per million.
